# Non-clinical safety profile and pharmacodynamics of two formulations of the anti-sepsis drug candidate Rejuveinix

**DOI:** 10.1101/2021.01.03.425139

**Authors:** Fatih M. Uckun, Cemal Orhan, Joy Powell, Emre Sahin, Ibrahim H. Ozercan, Michael Volk, Kazim Sahin

## Abstract

Here, we demonstrate that the two distinct formulations of our anti-sepsis drug candidate Rejuveinix (RJX) have a very favorable safety profile in Wistar Albino rats at dose levels comparable to the projected clinical dose levels. 14-day treatment with RJX-P or RJX-B similarly elevated the day 15 tissue levels of the antioxidant enzyme superoxide dismutase (SOD) as well as ascorbic acid in both the lungs and liver in a dose-dependent fashion. The activity of SOD and ascorbic acid levels were significantly higher in tissues of RJX-P or RJX-B treated rats than vehicle-treated control rats (*p* <0.0001). There was no statistically significant difference between tissues SOD activity or ascorbic acid levels of rats treated with RJX-P vs. rats treated with RJX-B (*p* >0.05). The observed elevations of the SOD and ascorbic acid levels were transient and were no longer detectable on day 28 following a 14-day recovery period. These results demonstrate that RJX-P and RJX-B are bioequivalent relative to their pharmacodynamic effects on tissue SOD and ascorbic acid levels.

## Introduction

Sepsis represents a strong inflammatory response to an infection with a potentially fatal outcome due to its complications, including acute respiratory distress syndrome (ARDS), septic shock, multiple organ dysfunction syndrome (MODS), and disseminated intravascular coagulopathy. Our anti-sepsis drug candidate, Rejuveinix (RJX), is a rationally-designed formulation of naturally occurring antioxidants and anti-inflammatory compounds, and it is being evaluated for its clinical impact potential for COVID-19 associated viral sepsis in a placebo-controlled, double-blind Phase II study RPI015 [1–3]. The composition, mode of action, and recently published favorable clinical safety profile of RJX [1–3] makes it an attractive anti-inflammatory drug candidate for the prevention and treatment of sepsis. We recently completed a Phase 1, double-blind, placebo-controlled, randomized, two-part, ascending dose-escalation study that evaluated its safety and tolerability in healthy volunteers (Protocol No. RPI003; ClinicalTrials.gov Identifier: NCT03680105) [3]. No deaths, serious adverse events (SAEs), or Grade 3-4 adverse events (AEs) were observed in any of 57 healthy volunteers treated with RJX at dose levels ranging from 0.024 mL/kg to 0.759 mL/kg [3].

The primary objective of the present non-clinical study was to compare the toxicity profiles and pharmacodynamics features of two formulations of RJX that are in clinical development using Wistar Albino rats. Our results presented herein demonstrate that the two formulations of RJX are bioequivalent relative to their pharmacodynamic effects on tissue SOD and ascorbic acid levels as well as safety profile in rodents.

## Materials and Methods

### RJX Formulations

Formulation RJX-P and RJX-B are two distinct intravenous formulations of RJX. Both formulations contain ascorbic acid, cyanocobalamin, thiamine hydrochloride, riboflavin 5’ phosphate, niacinamide, pyridoxine hydrochloride, and calcium D-pantothenate. Each formulation has a Part 1 (Vial A) and a Part 2 (Vial B). Vial A contains the active ingredients and Vial B is the buffer solution that is mixed with Vial A prior to administration. The composition of Vial A for RJX-P and RJX-B are detailed in Tables 1 and 2.

**Table 1.** Composition of Vial A.

| Formulation RJX-P |  |  |  |  |  |
| --- | --- | --- | --- | --- | --- |
| Test | Specification |  |  | Test Results |  |
| Appearance | Brownish-orange/reddish-orange |  |  | conforms |  |
| pH | 2.2 - 3.8 |  |  | 3.0 |  |
| Sterility | USP <71> |  |  | Sterile |  |
| Endotoxin | USP <85>, NMT 17.5 EU/mL – MVD = 1:3500 |  |  | <2.5000 EU/mL |  |
| Particulate Matter | USP <788>, The average number of particles present in the units tested does not exceed 6000 per container $\geq 10 \mu\text{m}$ and does not exceed 600 per container $\geq 25 \mu\text{m}$ | | | $\geq 10 \mu\text{m}$: 15 particles/container;<br>$\geq 25 \mu\text{m}$: 0 particles/container | |
| Assay | Target (% content by weight) | Lower Limit (% content by weight) | Upper Limit (% content by weight) | Measured % content | % of Specified Target Content |
| Ascorbic Acid | 8.9933 | 8.094 | 9.893 | 9.011 | 100.2 |
| Niacinamide | 1.188 | 1.069 | 1.307 | 1.186 | 99.8 |
| Pyridoxine HCl | 1.188 | 1.069 | 1.307 | 1.191 | 100.3 |
| Riboflavin 5' Phosphate | 0.0253 | 0.023 | 0.028 | 0.025 | 98.8 |
| Thiamine HCl | 0.6333 | 0.57 | 0.697 | 0.643 | 101.5 |
| Calcium Pantothenate | 0.0293 | 0.026 | 0.032 | 0.028 | 95.6 |
| Cyanocobalamin | 0.0193 | 0.017 | 0.021 | 0.019 | 98.4 |
| Magnesium Sulfate | 8.080 | 7.272 | 8.888 | 8.176 | 101.2 |

**Table 2.** Composition of Vial A.

| **Formulation RJX-B** |  |  |  |  |  |
| --- | --- | --- | --- | --- | --- |
| **Test** | **Specification** |  |  | **Test Results** |  |
| **Appearance** | Brownish-orange/reddish-orange |  |  | conforms |  |
| **pH** | 2.2 - 3.8 |  |  | 3.0 |  |
| **Sterility** | USP <71> |  |  | Sterile |  |
| **Endotoxin** | USP <85>, NMT 17.5 EU/mL – MVD = 1:3500 |  |  | <2.5000 EU/mL |  |
| **Particulate Matter** | USP <788>, The average number of particles present in the units tested does not exceed 6000 per container $\geq 10 \mu\text{m}$ and does not exceed 600 per container $\geq 25 \mu\text{m}$ | | | $\geq 10 \mu\text{m}$: 123 particles/container;<br>$\geq 25 \mu\text{m}$: 1 particle/container | |
| **Assay** | Target (% content by weight) | Lower Limit (% content by weight) | Upper Limit (% content by weight) | Measured % content | % of Specified Target Content |
| **Ascorbic Acid** | 7.6443 | 6.880 | 8.409 | 7.751 | 101.4 |
| **Niacinamide** | 1.010 | 0.909 | 1.111 | 1.017 | 100.7 |
| **Pyridoxine HCl** | 1.010 | 0.909 | 1.111 | 1.025 | 101.5 |
| **Riboflavin 5' Phosphate** | 0.0215 | 0.019 | 0.024 | 0.022 | 102.3 |
| **Thiamine HCl** | 0.5383 | 0.484 | 0.592 | 0.547 | 101.6 |
| **Calcium Pantothenate** | 0.0249 | 0.022 | 0.027 | 0.023 | 92.4 |
| **Cyanocobalamin** | 0.0164 | 0.015 | 0.018 | 0.017 | 103.7 |
| **Magnesium Sulfate** | 6.868 | 6.181 | 7.555 | 6.867 | 100.0 |

### Dose Preparation

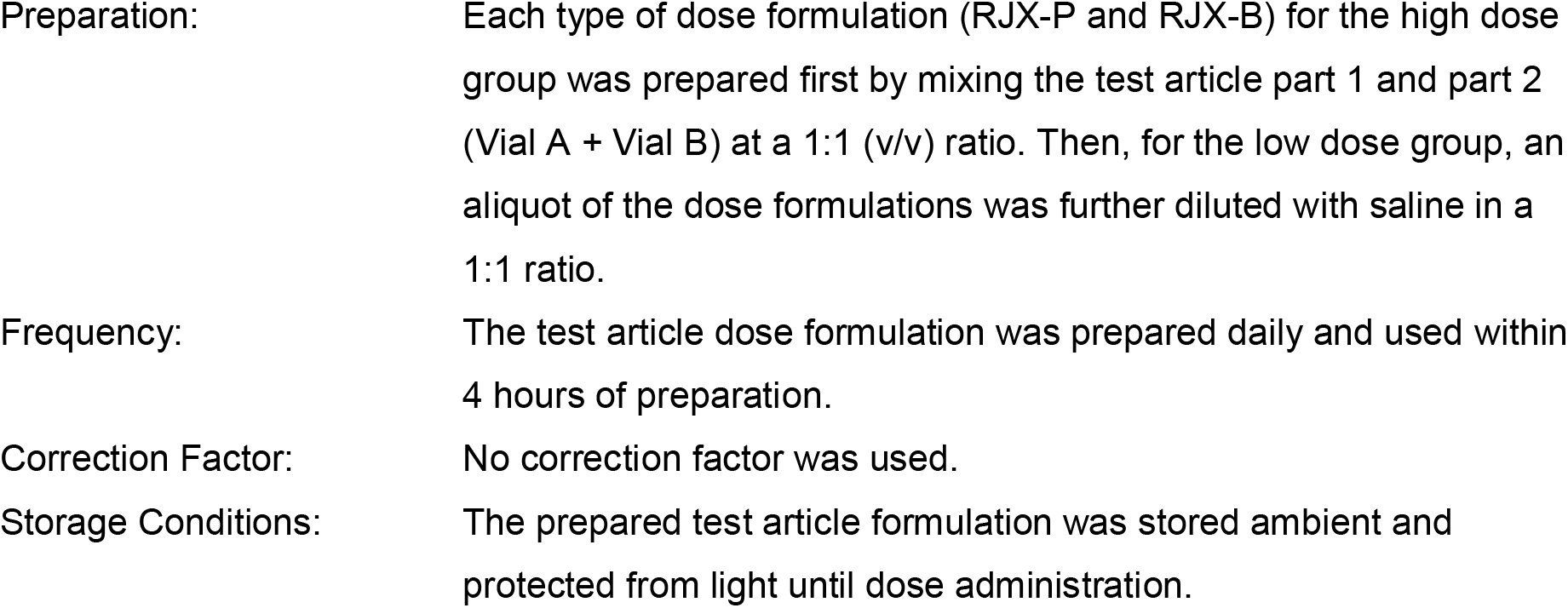
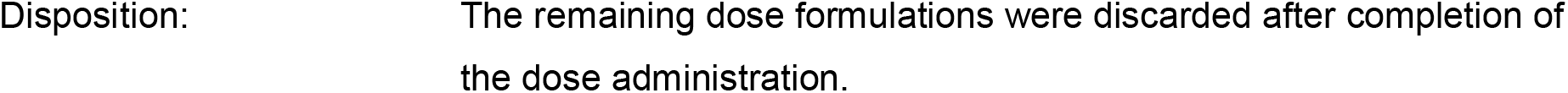

Table 5 compares the quantitative composition of RJX-P and RJX-B.

**Table 3.** Composition of Vial B.

| Formulation RJX-P |  |  |  |  |  |
| --- | --- | --- | --- | --- | --- |
| Test | Specification |  |  | Test Results |  |
| Appearance | Clear, colorless, free of particulate |  |  | conforms |  |
| pH | 6.2 – 8.4 |  |  | 8.3 |  |
| Sterility | USP <71> |  |  | Sterile |  |
| Endotoxin | USP <85>, NMT 17.5 EU/mL – MVD = 1:3500 |  |  | <0.0500 EU/mL |  |
| Particulate Matter | USP <788>, The average number of particles present in the units tested does not exceed 6000 per container $\geq 10 \mu\text{m}$ and does not exceed 600 per container $\geq 25 \mu\text{m}$ | | | $\geq 10 \mu\text{m}$: 10 particles/container;<br>$\geq 25 \mu\text{m}$: 3 particles/container | |
| Assay | Target (% content by weight) | Lower Limit (% content by weight) | Upper Limit (% content by weight) | Measured % content | % of Specified Target Content |
| Sodium Bicarbonate | 8.40 | 7.56 | 9.24 | 8.80 | 104.8 |

**Table 4.** Composition of Vial B.

| Formulation RJX-B |  |  |  |  |  |
| --- | --- | --- | --- | --- | --- |
| Test | Specification |  |  | Test Results |  |
| Appearance | Clear, colorless, free of particulate |  |  | conforms |  |
| pH | 6.2 – 8.4 |  |  | 8.2 |  |
| Sterility | USP <71> |  |  | Sterile |  |
| Endotoxin | USP <85>, NMT 17.5 EU/mL – MVD = 1:3500 |  |  | <0.0500 EU/mL |  |
| Particulate Matter | USP <788>, The average number of particles present in the units tested does not exceed 6000 per container $\geq 10 \mu\text{m}$ and does not exceed 600 per container $\geq 25 \mu\text{m}$ | | | $\geq 10 \mu\text{m}$: 59 particles/container;<br>$\geq 25 \mu\text{m}$: 2 particles/container | |
| Assay | Target (% content by weight) | Lower Limit (% content by weight) | Upper Limit (% content by weight) | Measured % content | % of Specified Target Content |
| Sodium Bicarbonate | 8.40 | 7.56 | 9.24 | 8.44 | 100.5 |

**Table 5.** Quantitative Composition of RJX-P and RJX-B.

|  | Actual Measured %<br>content by weight |  | mg/mL in Vial A |  | mg/mL in 1:1 A:B |  |
| --- | --- | --- | --- | --- | --- | --- |
| Lot Number | RJX<br>PPP.18.1051 | RJX-B<br>200702-CLN | RJX<br>PPP.18.1051 | RJX-B<br>200702-CLN | RJX<br>PPP.18.1051 | RJX-B<br>200702-CLN |
| Ascorbic Acid | 9.011 | 7.751 | 90.11 | 77.51 | 45.055 | 38.755 |
| Thiamine | 0.643 | 0.547 | 6.43 | 5.47 | 3.215 | 2.735 |
| Magnesium Sulfate | 8.176 | 6.867 | 81.76 | 68.67 | 40.88 | 34.335 |
| Cyanocobalamin | 0.019 | 0.017 | 0.19 | 0.17 | 0.095 | 0.085 |
| Niacinamide | 1.186 | 1.017 | 11.86 | 10.17 | 5.93 | 5.085 |
| Pyridoxine | 1.191 | 1.025 | 11.91 | 10.25 | 5.955 | 5.125 |
| Riboflavin<br>5'Phosphate | 0.025 | 0.022 | 0.25 | 0.22 | 0.125 | 0.11 |
| Calcium D-Pantothenate | 0.028 | 0.023 | 0.28 | 0.23 | 0.14 | 0.115 |

### Animal Study

The toxicity of RJX-P and RJX-B formulations were examined at the Bingol University Experimental Animal Center in Bingol, Turkey with the approval of the Animal Care and Use Committee (IACUC) at the Bingol University (23092020-E-2310). 8-week old Wistar albino rats were used in this study. A total of 100, 20 rats per group were used. There were 2 cohorts. One cohort was treated for 14 days and sacrificed on day 15. The second cohort was treated for 14 days and sacrificed on day 28 following a 2 week recovery period. In each cohort, there were 5 treatment groups, including one control group and 4 RJX treatment groups. The collective group designation for cohort 1 + cohort 2 (N=100) was as follows:

**Group 1** (20 Rats, 10 female, 10 male; Cohort 1 = 5 female, 5 male; Cohort 2 = 5 female, 5 male): Vehicle control = 0.5 mL normal saline (NS)
**Group 2** (20 Rats, 10 female, 10 male; Cohort 1 = 5 female, 5 male; Cohort 2 = 5 female, 5 male): RJX-P - 0.5 mL × 14 days
**Group 3** (20 Rats,10 female, 10 male; Cohort 1 = 5 female, 5 male; Cohort 2 = 5 female, 5 male): RJX-P – 0.25 mL × 14 days.
**Group 4** (20 Rats, 10 female, 10 male; Cohort 1 = 5 female, 5 male; Cohort 2 = 5 female, 5 male): RJX-B – 0.5 mL RJX × 14 days
**Group 5** (20 Rats, 10 female, 10 male; Cohort 1 = 5 female, 5 male; Cohort 2 = 5 female, 5 male): RJX-B – 0.25 mL RJX × 14 days

Rats in Cohort 1 (5 males and 5 females from each group listed above) were observed for clinical signs of toxicity daily, weighed biweekly, and sacrificed on day 15 or day 28 to determine the toxicity of RJX by examining their blood chemistry profiles, blood counts, and histopathological examinations of tissue specimens. Blood was collected after decapitation by cervical dislocation. All laboratory analyses were performed at the Firat University (Elazig, Turkey). The blood chemistry profiles were examined using automated laboratory methods (Samsung Labgeo PT10, Suwon, Korea). Blood counts (red blood cells [RBC], white blood cells [WBC], and platelets [Plt]) were determined using an Exigo Hematology Analyzer (Sweden).

Blood samples were used for safety pharmacology and serum chemistry tests. Serum concentrations of aspartate aminotransferase (AST), alanine aminotransferase (ALT), alkaline phosphatase (ALP) and total bilirubin, were assayed using an automated chemistry analyzer (Samsung LABGEO PT10, Samsung Electronics Co., Suwon, Korea). Safety pharmacology labs, tissue activity of superoxide dismutase (SOD), and tissue ascorbic acid levels were determined to examine the effects of RJX-P and RJX-B. The activities of superoxide dismutase (SOD) were determined by quantitative enzyme-linked immunosorbent assays (ELISA, Bio-Tek Elx800 Universal Microplate Reader, Bio-Tek Instruments, Inc, Winooski, USA) using commercially available kits (Cayman Chemical, Ann Arbor, MI, USA) [3]. At the time of necropsy, multiple tissues (Liver, Lung, Kidney, Spleen, Heart, Pancreas, Brain, Cerebellum, Intestine, Stomach, Testes, Uterus, Skeletal Muscle, and Skin) were collected within 15 min after sacrifice for gross pathological and histopathological examinations. Organs were preserved in 10% neutral phosphate-buffered formalin and processed for histologic sectioning. For histopathologic studies, formalin-fixed tissues were dehydrated and embedded in paraffin by routine methods. Glass slides with affixed 4-5 micron tissue sections were prepared and stained with hematoxylin and Eosin (H&E). Sections were examined by light microscopy with an Olympus BX-50 microscope (Center Valley, PA).

### Test System Justification

The rat is a standard rodent species for general toxicity study. Intraperitoneal dosing achieves the expected systemic exposure of the drug for toxicity evaluation. The number of animals per group (5/sex/group) was the minimal number for each cohort that allows for sufficient data point collection and statistical analysis of several endpoints for the determination of systemic effects.

### Details of In-Life Procedures

#### Experimental Design for Each Cohort

| Group | Dose Assignment | Number of Animals | Dose Conc. (mg/mL) | Dose Level (mL/kg/day) | Dose Volume (mL/animal/day) |
| --- | --- | --- | --- | --- | --- |
| 1 | Control (Saline) | 5 M/5 F | 0 | 0 | 0.5 |
| 2 | RJX-P - High Dose | 5 M/5 F | * | 2.5 | 0.5 (0.5 mL RJX-P) |
| 3 | RJX-P - Low Dose | 5 M/5 F | * | 1.25 | 0.5 (0.25 mL RJX-P +0.25 mL NS) |
| 4 | RJX-B - High Dose | 5 M/5 F | * | 2.5 | 0.5 (0.5 mL RJX-B) |
| 5 | RJX-B - Low Dose | 5 M/5 F | * | 1.25 | 0.5 (0.25 mL RJX-B +0.25 mL NS) |

M = male; F = female; Conc. = concentration
\* The test article is a solution formulated with multiple components of varying content concentration levels, at fixed ratios. See Table 1 for details of target formulation content.

### Dose Level Justification

The purpose of the rat toxicity study is to compare the toxicity of two RJX formulations to rule out unexpected, unusual severe toxicities at the clinically applicable dose levels. The toxicity of RJX in rodents has been extensively studied. The no observed adverse effect level (NOAEL) was >5 mL/kg in both Sprague Dawley rats (when administered via intra-arterial route daily for 21 days) and Beagle dogs (when administered via intra-arterial route daily × 28 days). The HED of the >5mL/kg NOAEL in rats is 0.8 mL/kg and the HED of the ≥5 mL/kg NOAEL in dogs is 2.7 mL/kg. The safety of RJX in human subjects was examined in a Phase 1 study in healthy human volunteers and the MTD was determined to be 0.5 mL/kg. In a clinical study in COVID-19 patients (RPI015), RJX is being used at a fixed dose level of 20 mL/dose/patient. This corresponds to an estimated average per kg dose level of 0.25 mL/kg to 0.33 mL/kg for an adult patient weighing 60-80 kg. The selected higher dose level in rats is 2.5 mL/kg, which corresponds to a human equivalent dose of 2.5 × 0.16 mL/kg = 0.4 mL/kg. The dose is selected as a dose level ~20% higher than the estimated average clinical dose.

The selected lower dose level in rats is 1.25 mL/kg, which corresponds to a human equivalent dose of 0.2 mL/kg. This dose was selected as a dose level ~20% lower than the estimated low dose level in the clinical study for an 80 kg adult patient. Both dose levels are substantially higher than the 0.07 mL/kg (i.e., 0.42 mL/kg of a 1:6 diluted solution) effective dose level at which RJX was shown to both prevent and reverse cytokine storm, acute lung injury, and multi-organ dysfunction in a model of LPS-galactosamine induced sepsis, septic shock, ARDS, and multi-organ failure [18-uckun Frontiers-2020]. The same dose levels were selected for the testing of RJX-B for comparison purposes. As the concentration of the active ingredients is lower in RJX-B, the safety profile of this new formulation was postulated to be similar or better than the safety profile of RJX-P.

### Statistical Analysis

Statistical analysis includes analysis of variance (ANOVA) and/or, nonparametric analysis of variance (Kruskal-Wallis) using the SPSS statistical program (IBM, SPPS Version 21). Shapiro-Wilk test was performed for normality with a significance level of 0.05. If the data shows normally distribution (Shapiro-Wilk test result p ≥ 0.05), a parametric analysis of variance (ANOVA) was performed. After that, Levene’s test was performed for homogeneity with a significance level of 0.05. For homogeneous data (Levene’s test result p ≥ 0.05), a parametric analysis of variance (ANOVA) was performed and Tukey’s multiple comparisons were used as a post hoc test to detect alterations among the groups. If the data is not homogeneous (Levene’s test result p < 0.05), Dunnett’s tests (Dunnett’s T3 or Dunnett’s C test) were performed to compare treatment groups values to those of the vehicle/control group(s). If the ANOVA results were “not significantly different” (F test p ≥ 0.05), the analysis stopped. If the data did not show normally distribution (Shapiro-Wilk test result p < 0.05), a non-parametric analysis of variance (Kruskal-Wallis) was performed. If the Kruskal-Wallis test returned “statistically different” (p < 0.05), the Dunn’s multiple corporation test was performed to compare the values of the treatment groups to those of the vehicle/control group(s). If the Kruskal-Wallis test returned “not significantly different” (p ≥ 0.05), the analysis stopped. P-values <0.05 were considered significant.

## Results

### Toxicity

RJX-P and RJX-B, when administered as daily intraperitoneal bolus injections for 14 consecutive days with or without a 14-day recovery time were not toxic to female or male Wistar Albino rats at 0.25 mL or 0.5 mL dose levels. None of the 40 rats treated with RJX-P or 40 rats treated with RJX-B showed any signs of morbidity. Groups of 20 rats were sacrificed on day 15 and day 28 (14 day recovery period group), respectively. The majority of the RJX-P or RJX-B-treated rats, gained weight during the 0-14 day, 14-28 day and, 0-28 day observation period. The average weight gain was 4.20 g (2.1%) for vehicle-treated control rats, and 4.59 g (2.3%) and 4.55 g (2.3%) for rats treated with the higher dose of RJX-P or RJX-B for a 0-28 day period (**Table 6** and **Figure 1**). As shown in **Table 7** and **Table 8**, the blood chemistry and hematology profiles of these rats did not suggest any system-specific toxicity. In particular, RJX-P or RJX-B-treated rats showed: (*a*) normal serum BUN and creatinine levels consistent with absence of renal toxicity; and (*b*) normal WBC, ANC, ALC, and RBC counts consistent with absence of hematologic toxicity. Histopathological examination of multiple tissues from RJX-P or RJX-B -treated rats in the day 15 group (*n* = 20) (**Table 9**, **Figures 2–6**) or day 28 groups (n=10) (**Table 10**) did not reveal any toxic lesions․. Thus, RJX-P and RJX-B were not toxic to rats at dose levels up to 0.5 mL/kg.

**Table 6.** Effects of treatment with RJX-P or RJX-B intraperitoneal injections on body weight (in grams) of Wistar albino rats.

| Time | Control<br>(NS) | RJX-P<br>(0.5 mL) | RJX-P<br>(0.25 mL) | RJX-B<br>(0.5 mL) | RJX-B<br>(0.25 mL) |
| --- | --- | --- | --- | --- | --- |
| Day 1 | 201.15±21.64 | 201.40±20.59 | 200.65±20.04 | 201.35±20.81 | 201.45±22.34 |
| Day 14 | 230.30±24.48 | 234.65±20.52 | 232.95±25.75 | 231.70±24.68 | 232.05±23.40 |
| Day 28 | 259.90±39.77 | 265.70±42.77 | 261.00±49.76 | 265.10±49.62 | 261.20±44.64 |

**Table 7.** Effects of 14-day treatment with RJX-P or RJX-B intraperitoneal injections on safety pharmacology laboratory parameters of Wistar albino rats.

| Parameters | Control<br>(NS) | RJX-P<br>(0.5 mL) | RJX-P<br>(0.25 mL) | RJX-B<br>(0.5 mL) | RJX-B<br>(0.25 mL) |
| --- | --- | --- | --- | --- | --- |
| Glucose, mg/dL | 119.80±8.13 | 117.90±9.09 | 116.80±7.19 | 118.20±8.46 | 116.90±7.06 |
| BUN, mg/dL | 21.40±2.93 | 22.74±1.82 | 23.00±2.19 | 23.36±3.06 | 22.20±1.65 |
| TP, g/dL | 7.17±0.70 | 7.68±0.23 | 7.49±0.81 | 7.60±0.41 | 7.49±0.24 |
| ALB, g/dL | 3.76±0.37 | 3.68±0.15 | 3.65±0.38 | 3.68±0.27 | 3.59±0.23 |
| GLOB, g/dL | 3.72±0.37 | 3.84±0.39 | 3.77±0.54 | 3.81±0.23 | 3.74±0.18 |
| ALT, U/L | 125.50±14.49 | 133.30±11.20 | 131.50±5.44 | 130.50±15.80 | 135.20±10.85 |
| AST, U/L | 144.50±21.82 | 162.60±15.58 | 150.10±19.06 | 154.80±42.86 | 164.60±26.71 |
| ALP, U/L | 261.80±28.86 | 270.50±37.61 | 263.90±21.26 | 267.20±30.19 | 242.30±37.29 |
| TBIL, mg/dL | 0.23±0.05 | 0.21±0.01 | 0.23±0.05 | 0.25±0.03 | 0.22±0.04 |
| Creatinine,<br>mg/dL | 0.42 ±0.04 | 0.43±0.04 | 0.41±0.03 | 0.44±0.04 | 0.44±0.02 |
| WBC(x10<sup>3</sup>/μL) | 10.85±4.07 | 9.77±2.62 | 12.37±3.14 | 11.33±2.22 | 10.13±1.24 |
| ALC(x10<sup>3</sup>/μL) | 7.56±2.89 | 6.96±2.07 | 8.90±2.57 | 7.86±1.17 | 7.09±0.95 |
| MON#(x10<sup>3</sup>/μL) | 0.67±0.32 | 0.58±0.23 | 0.75±0.13 | 0.56±0.07 | 0.57±0.21 |
| ANC (x10<sup>3</sup>/μL) | 2.19±1.12 | 1.87±0.49 | 2.31±0.81 | 2.47±1.28 | 1.95±1.09 |
| EOS#(x10<sup>3</sup>/μL) | 0.43±0.30 | 0.36±0.20 | 0.41±0.24 | 0.44±0.14 | 0.52±0.31 |
| LYM% | 70.08±6.98 | 70.85±3.27 | 71.91±5.14 | 70.07±6.41 | 70.48±8.84 |
| MON% | 5.69±1.24 | 5.45±1.55 | 5.81±0.98 | 4.78±0.58 | 5.24±1.37 |
| NEU% | 20.44±5.75 | 19.76±3.28 | 18.97±4.73 | 21.30±6.86 | 19.14±8.71 |
| EOS% | 3.79±2.80 | 3.94±2.01 | 3.31±1.62 | 3.85±1.10 | 5.14±2.96 |
| RBC (x10<sup>6</sup>/μL) | 7.83±0.50 | 8.03±0.81 | 7.69±1.06 | 7.83±1.41 | 8.25±0.40 |
| HGB, g/dL | 16.72±1.16 | 16.13±3.17 | 16.17±2.06 | 16.61±2.85 | 17.36±0.73 |
| HCT, % | 48.49±3.89 | 49.27±3.32 | 47.40±3.07 | 46.21±8.11 | 50.93±4.90 |
| MCV, fL | 62.04±5.38 | 61.69±4.76 | 62.46±7.34 | 59.07±2.35 | 61.94±7.35 |
| MCH, pg | 21.37±1.01 | 21.40±1.16 | 21.10±0.86 | 21.25±0.56 | 21.07±0.83 |
| MCHC, g/Dl | 34.63±2.45 | 34.79±1.45 | 34.06±3.23 | 35.99±1.00 | 34.39±3.66 |
| RDWa, fL | 43.84±7.29 | 44.47±7.11 | 48.49±10.18 | 44.60±6.55 | 45.56±10.13 |
| RDW, % | 16.63±1.05 | 17.05±1.28 | 17.55±1.08 | 16.42±0.46 | 16.47±1.07 |
| PLT(x10<sup>3</sup>/μL) | 723.10±124.39 | 802.50±170.25 | 787.20±159.14 | 722.40±127.12 | 723.10±80.93 |
| MPV, fL | 6.76±0.33 | 6.56±0.14 | 6.55±0.47 | 6.76±0.44 | 6.66±0.39 |

**Table 8.**
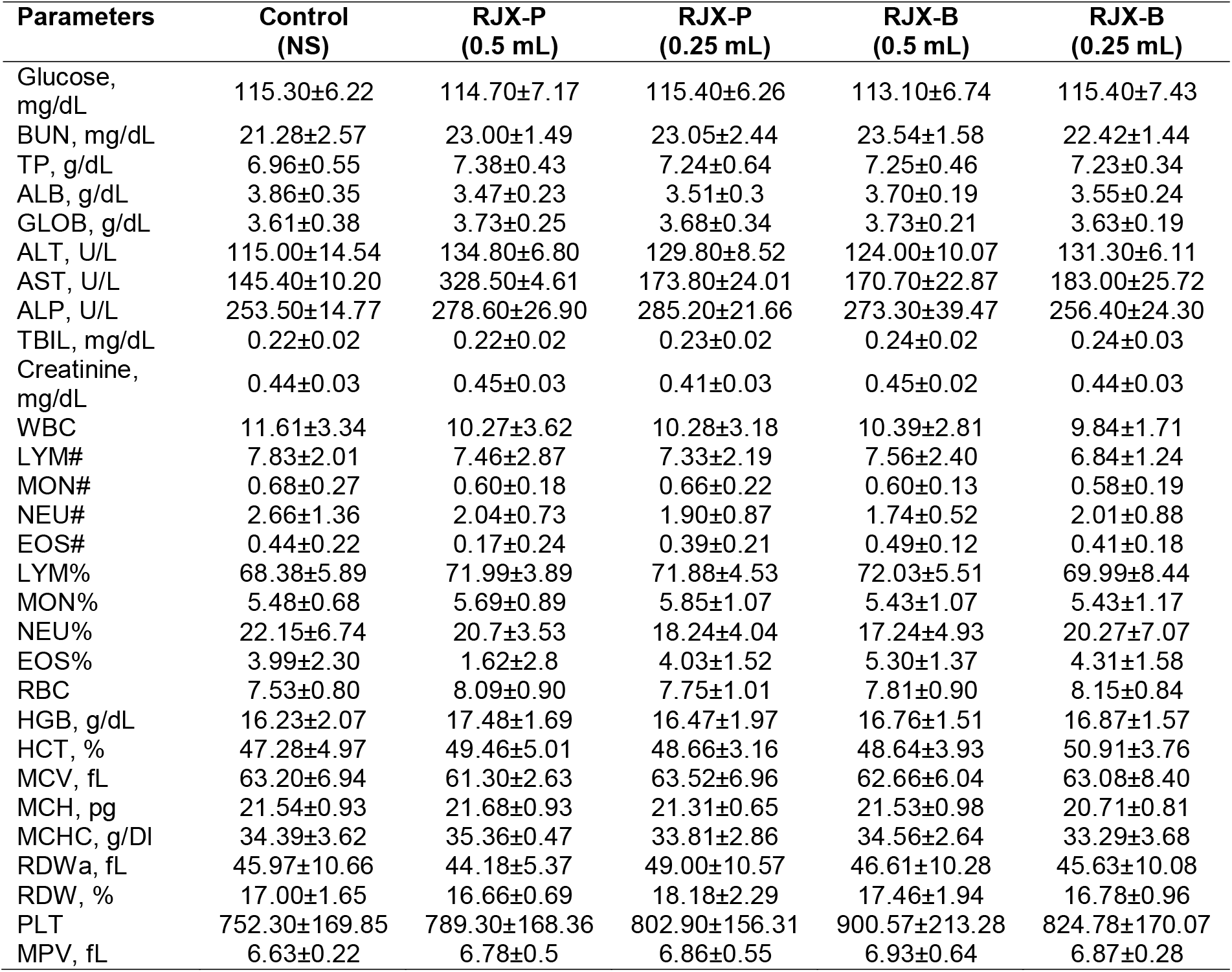
Effects of 14-day treatment with RJX-Por RJX-B intraperitoneal injections followed by 14 days recovery period on safety pharmacology laboratory parameters of Wistar albino rats.

**Table 9.**
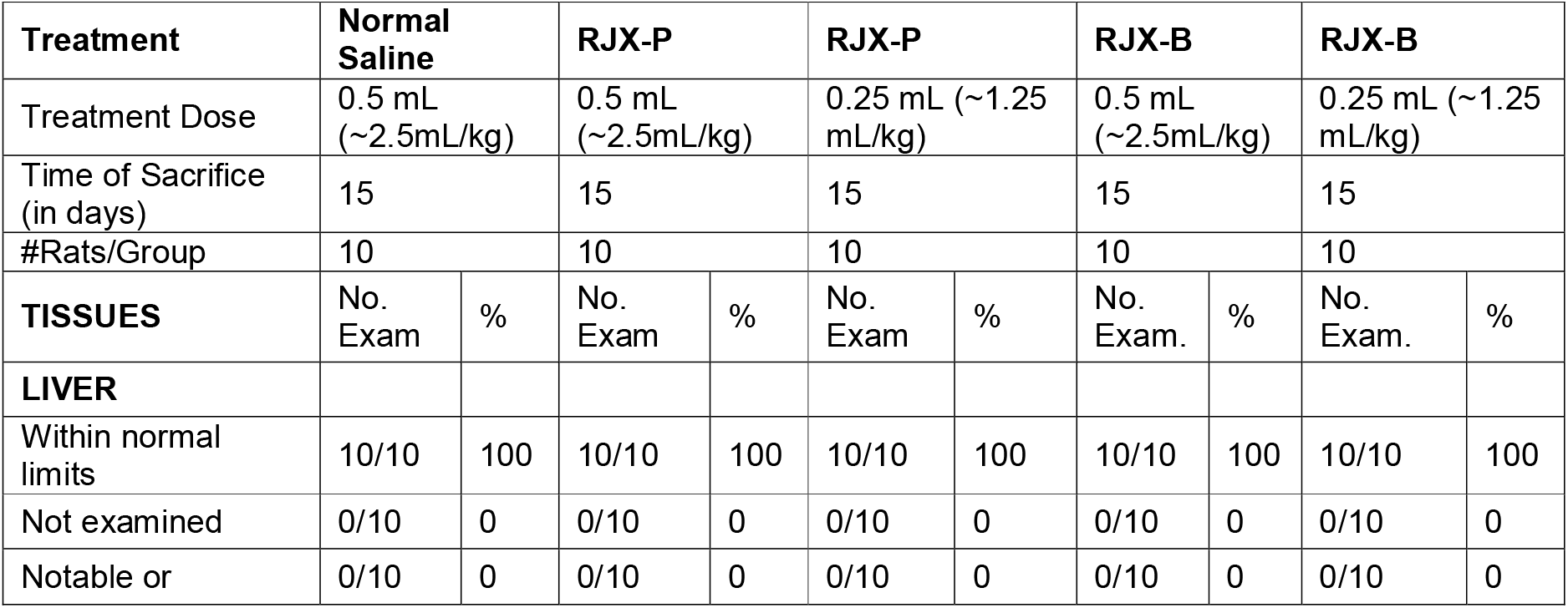
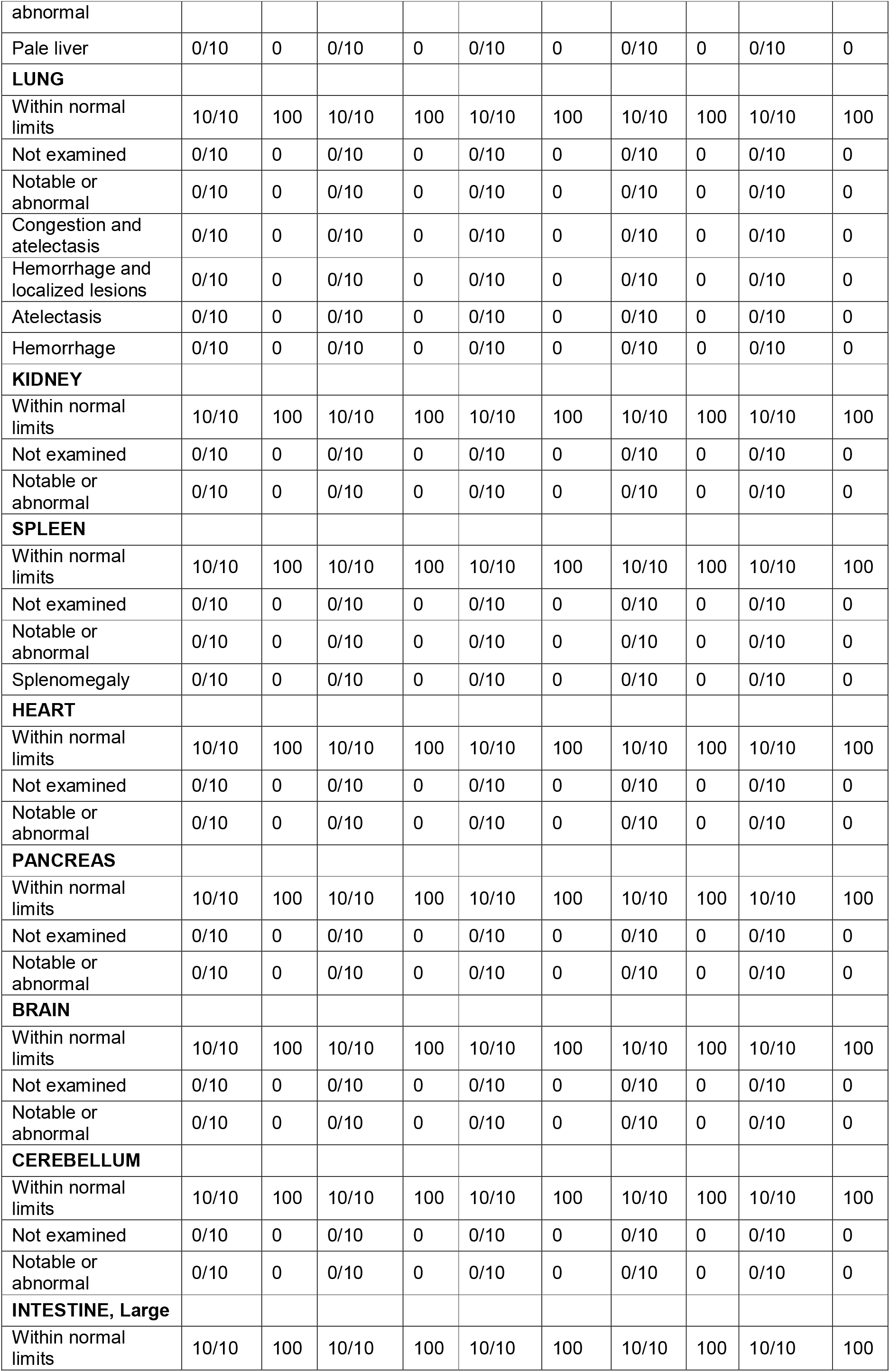
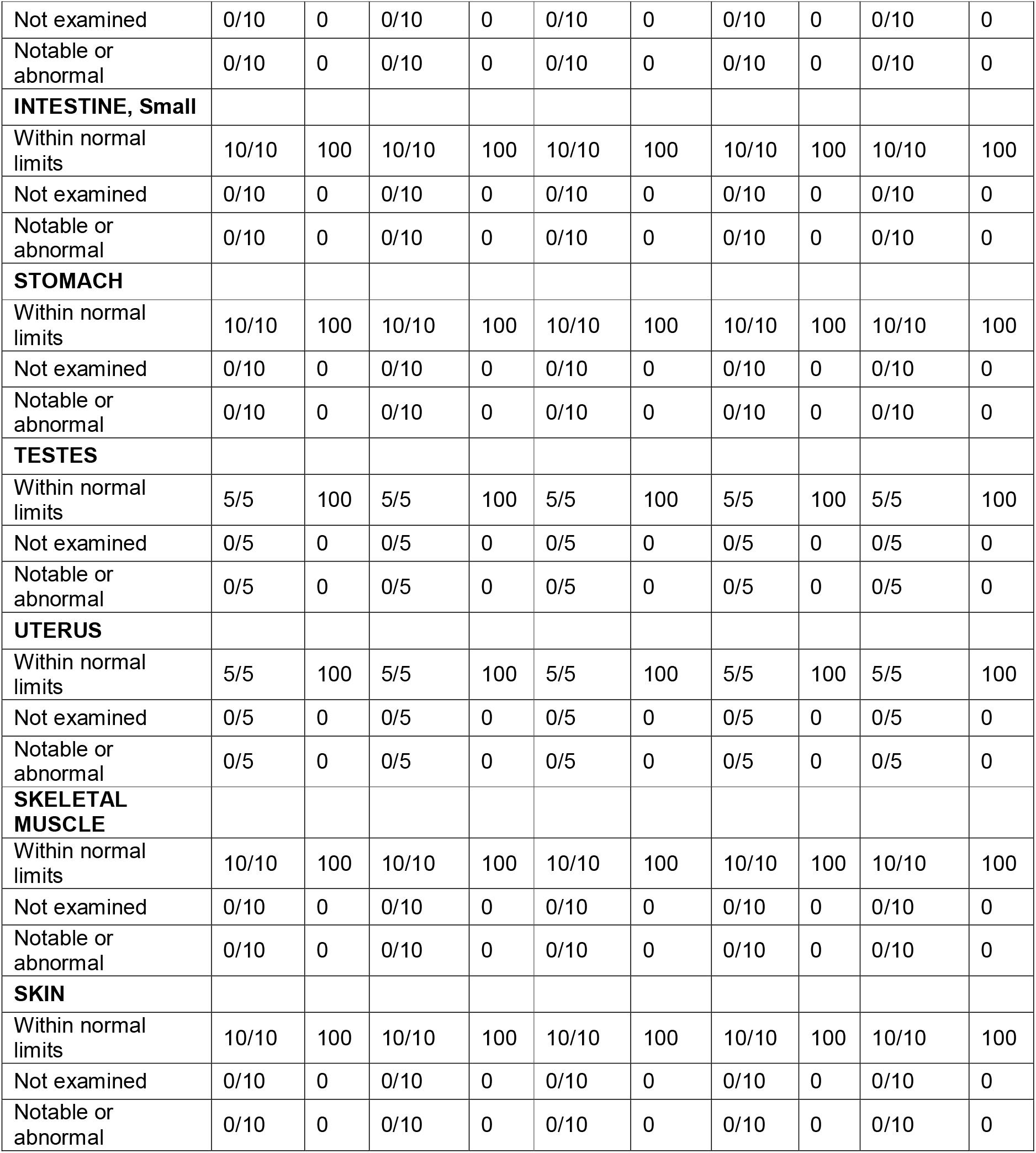
Histopathology findings in Wistar albino rats treated with 14 days of RJX-P or RJX-B intraperitoneal injections. Use summary table.

**Table 10.**
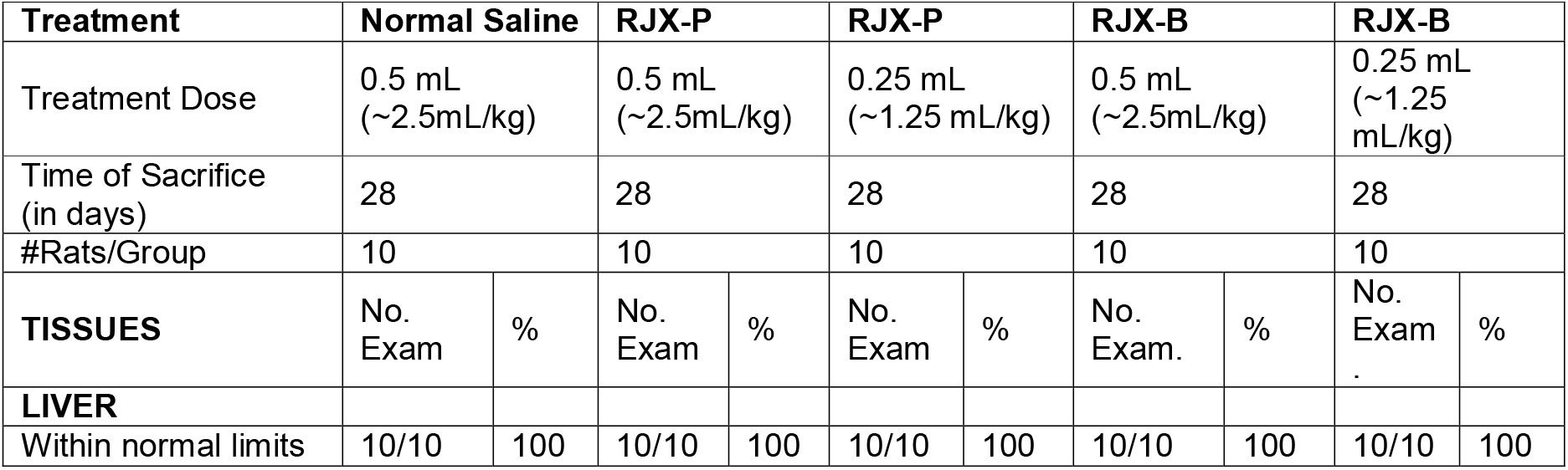
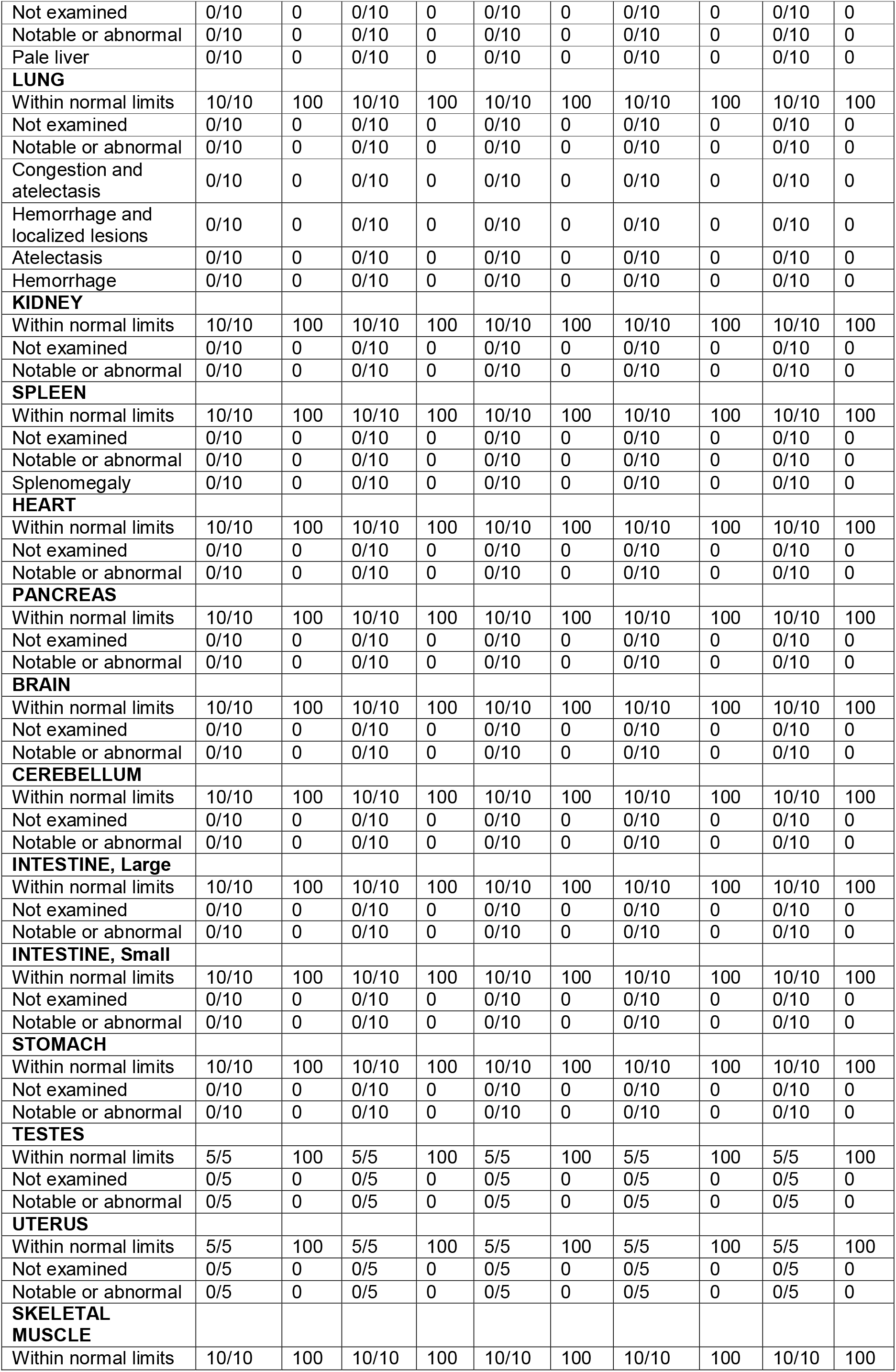
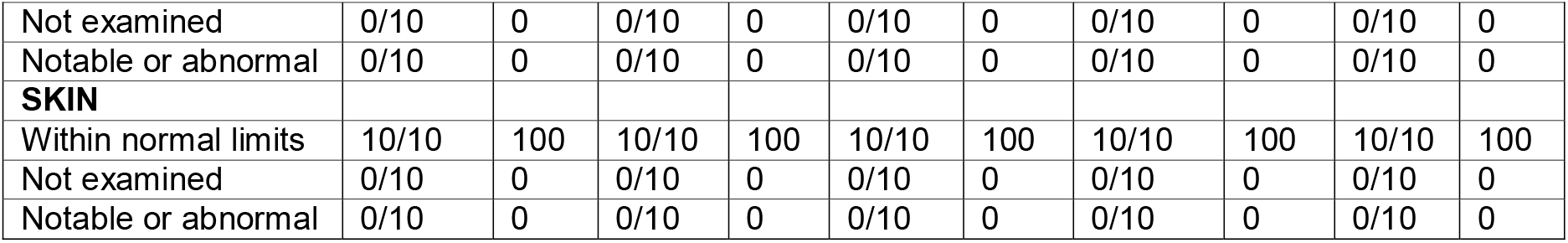
Day 28 Histopathology findings in Wistar albino rats treated with 14 days of RJX-P or RJXB intraperitoneal injections followed by 14 days recovery period.

**Figure 1.**
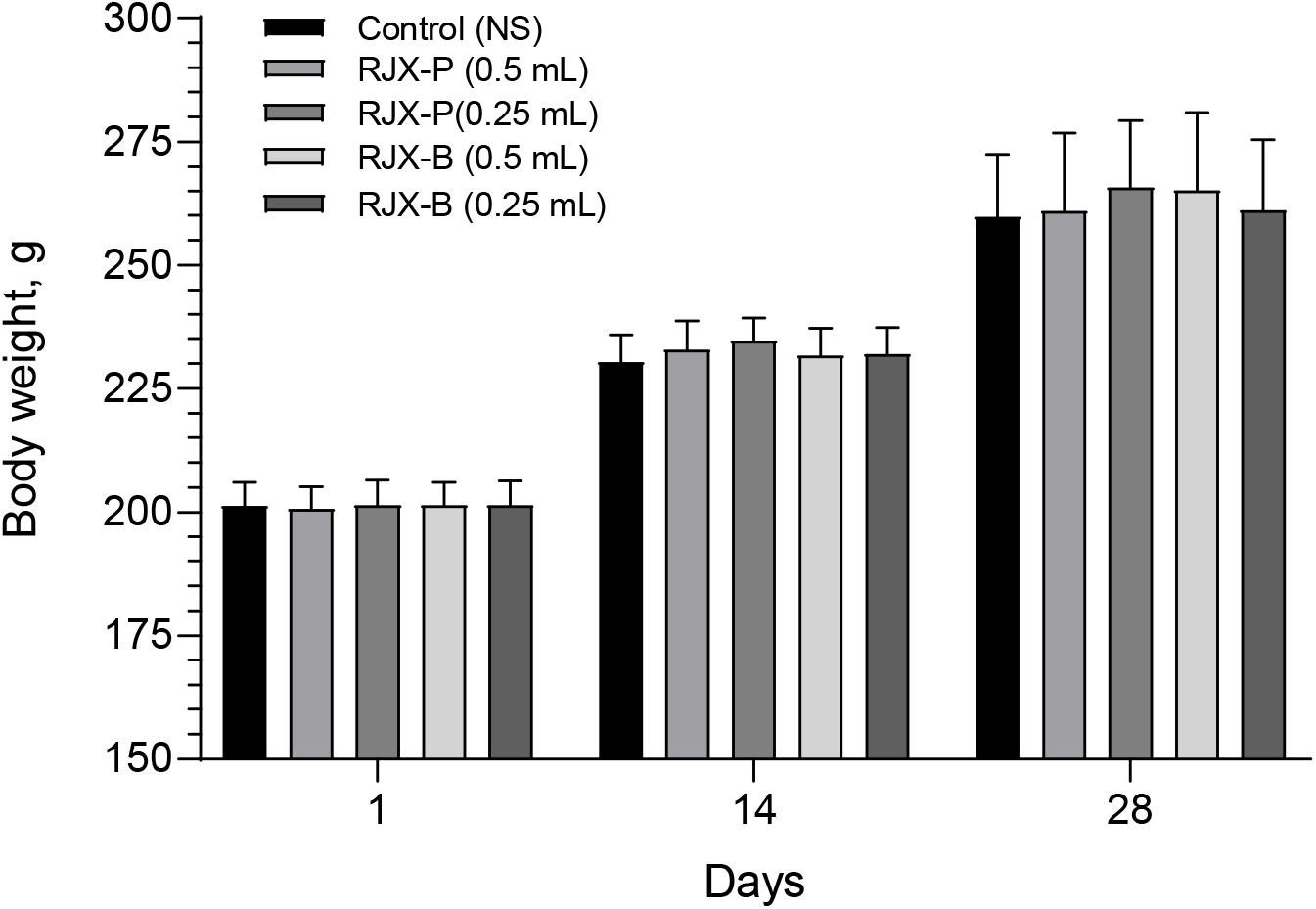
Effects of treatment with RJX-P or RJX-B intraperitoneal injections on body weight of Wistar albino rats.

**Figure 2.**
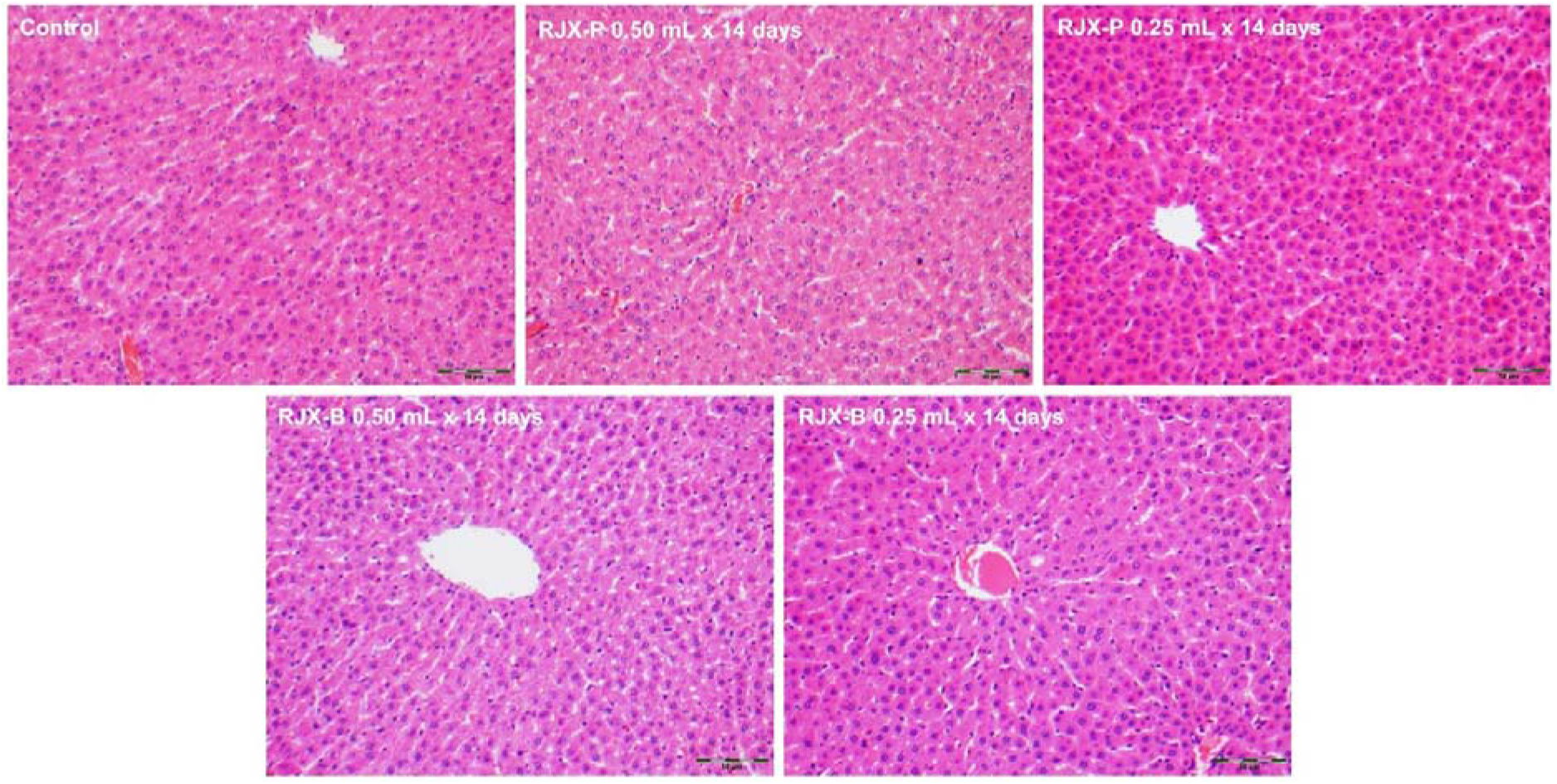
The day-15 histopathology of liver tissue in rats treated with 14 days of RJX-P or RJX-B intraperitoneal injections. Groups of 10 Wistar albino rats were treated with i.p injections of RJX-P (1.25 and 2.50 mL/kg/day) or RJX-B (1.25 and 2.50 mL/kg/day), or vehicle (NS) for 14 days. Animals were sacrificed on day 15 for histopathologic examinations. No toxic lesions were detected. H&E, 200x.

**Figure 3.**
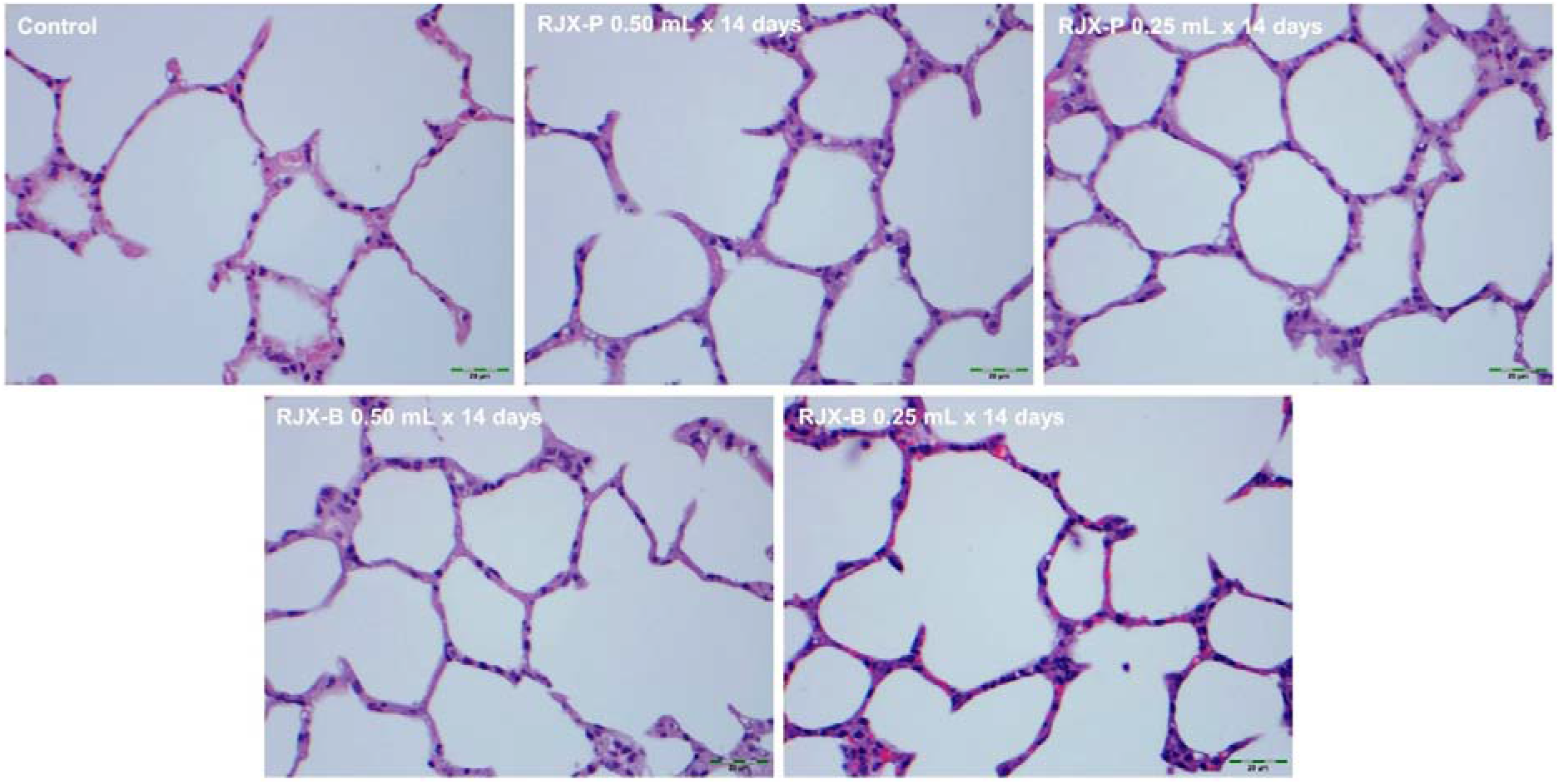
The day-15 histopathology of lung tissue in rats treated with 14 days of RJX-P or RJX-B intraperitoneal injections. Groups of 10 Wistar albino rats were treated with i.p injections of RJX-P (1.25 and 2.50 mL/kg/day) or RJX-B (1.25 and 2.50 mL/kg/day), or vehicle (NS) for 14 days. Animals were sacrificed on day 15 for histopathologic examinations. No toxic lesions were detected. H&E, 400×.

**Figure 4.**
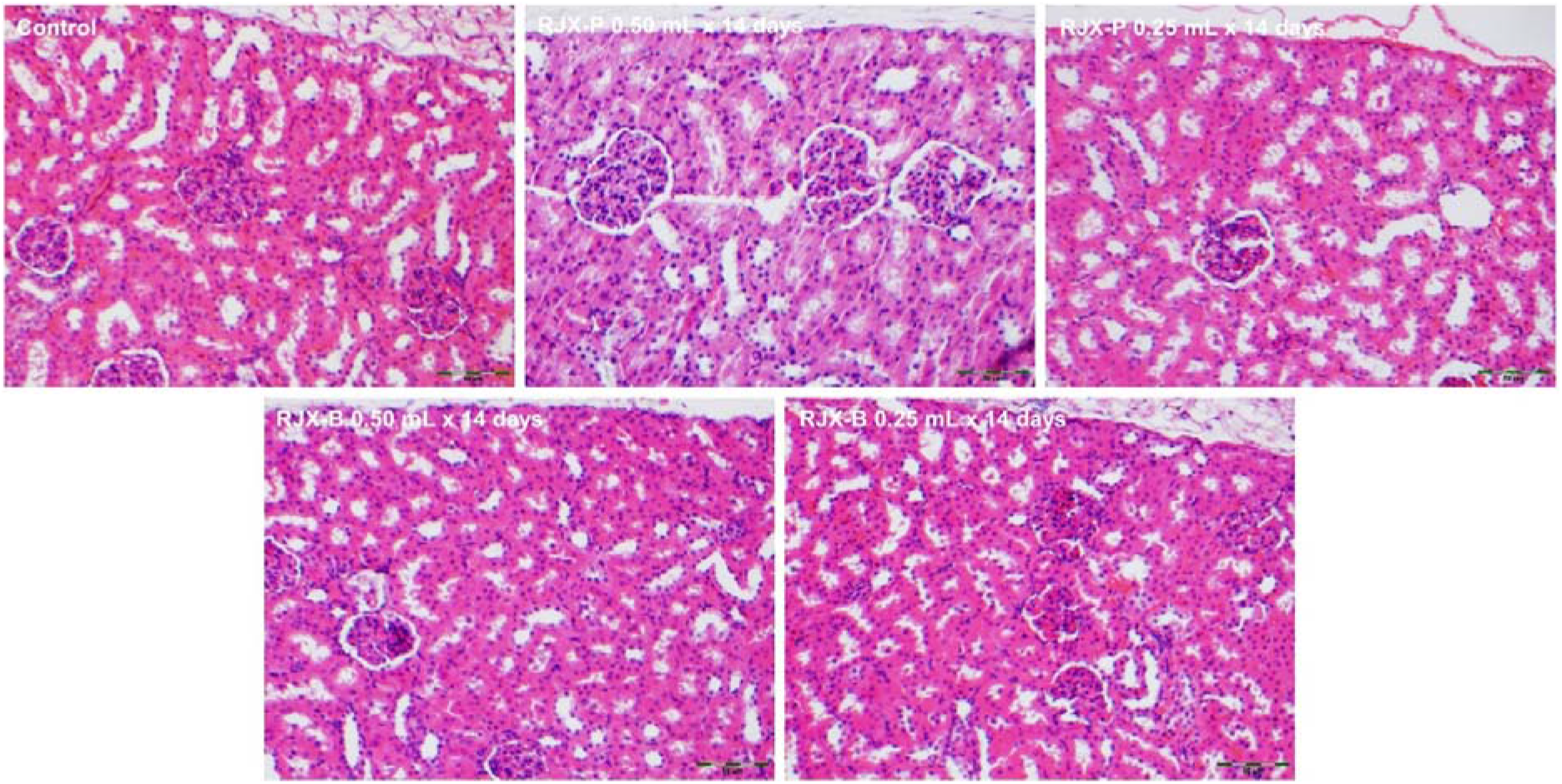
The day-15 histopathology of kidney tissue in rats treated with 14 days of RJX-P or RJX-B intraperitoneal injections. Groups of 10 Wistar albino rats were treated with i.p injections of RJX-P (1.25 and 2.50 mL/kg/day) or RJX-B (1.25 and 2.50 mL/kg/day), or vehicle (NS) for 14 days. Animals were sacrificed on day 15 for histopathologic examinations. No toxic lesions were detected. H&E, 200x.

**Figure 5.**
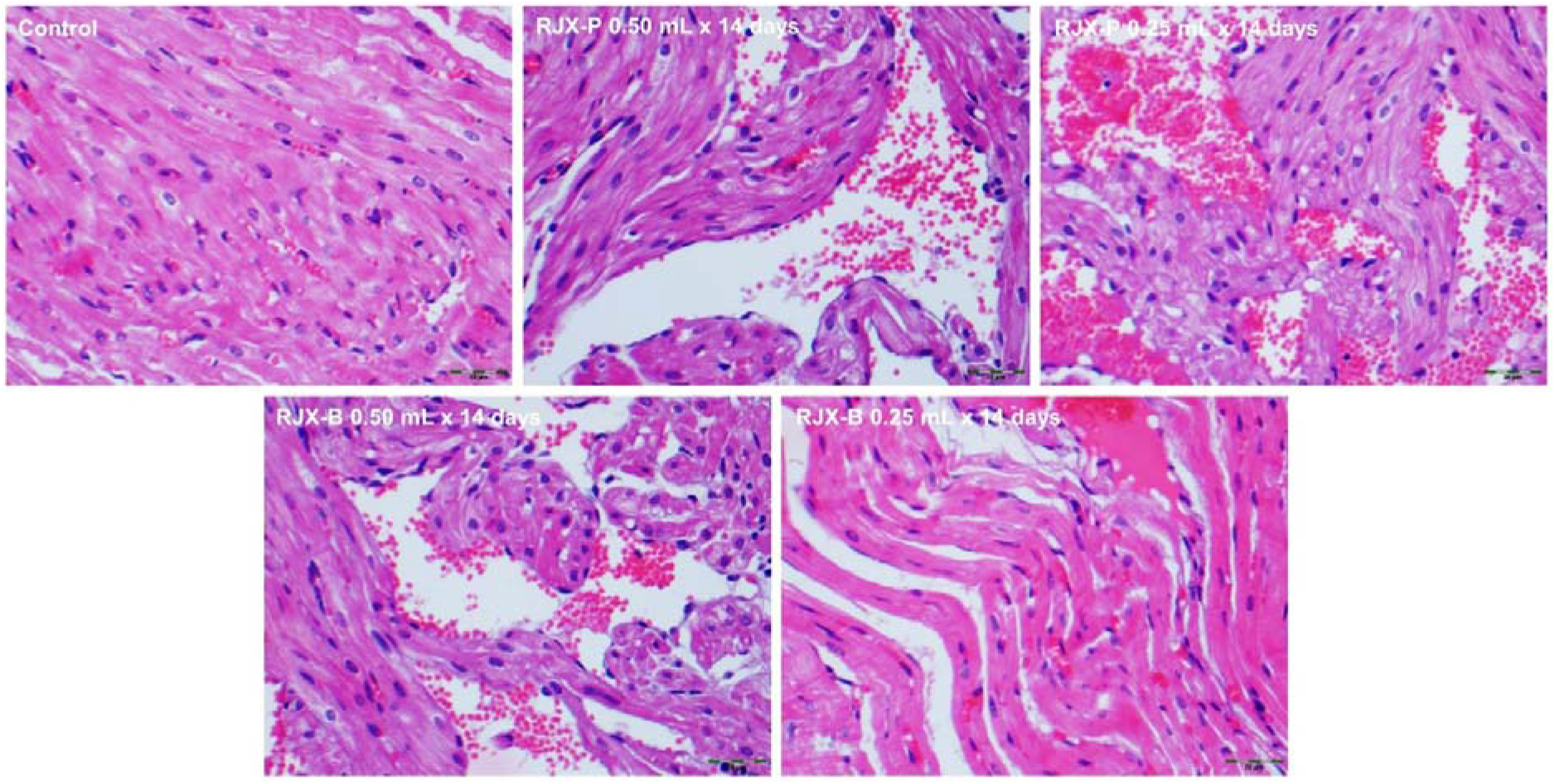
The day-15 histopathology of heart tissue in rats treated with 14 days of RJX-P or RJX-B intraperitoneal injections. Groups of 10 Wistar albino rats were treated with i.p injections of RJX-P (1.25 and 2.50 mL/kg/day) or RJX-B (1.25 and 2.50 mL/kg/day), or vehicle (NS) for 14 days. Animals were sacrificed on day 15 for histopathologic examinations. No toxic lesions were detected. H&E, 400×.

**Figure 6.**
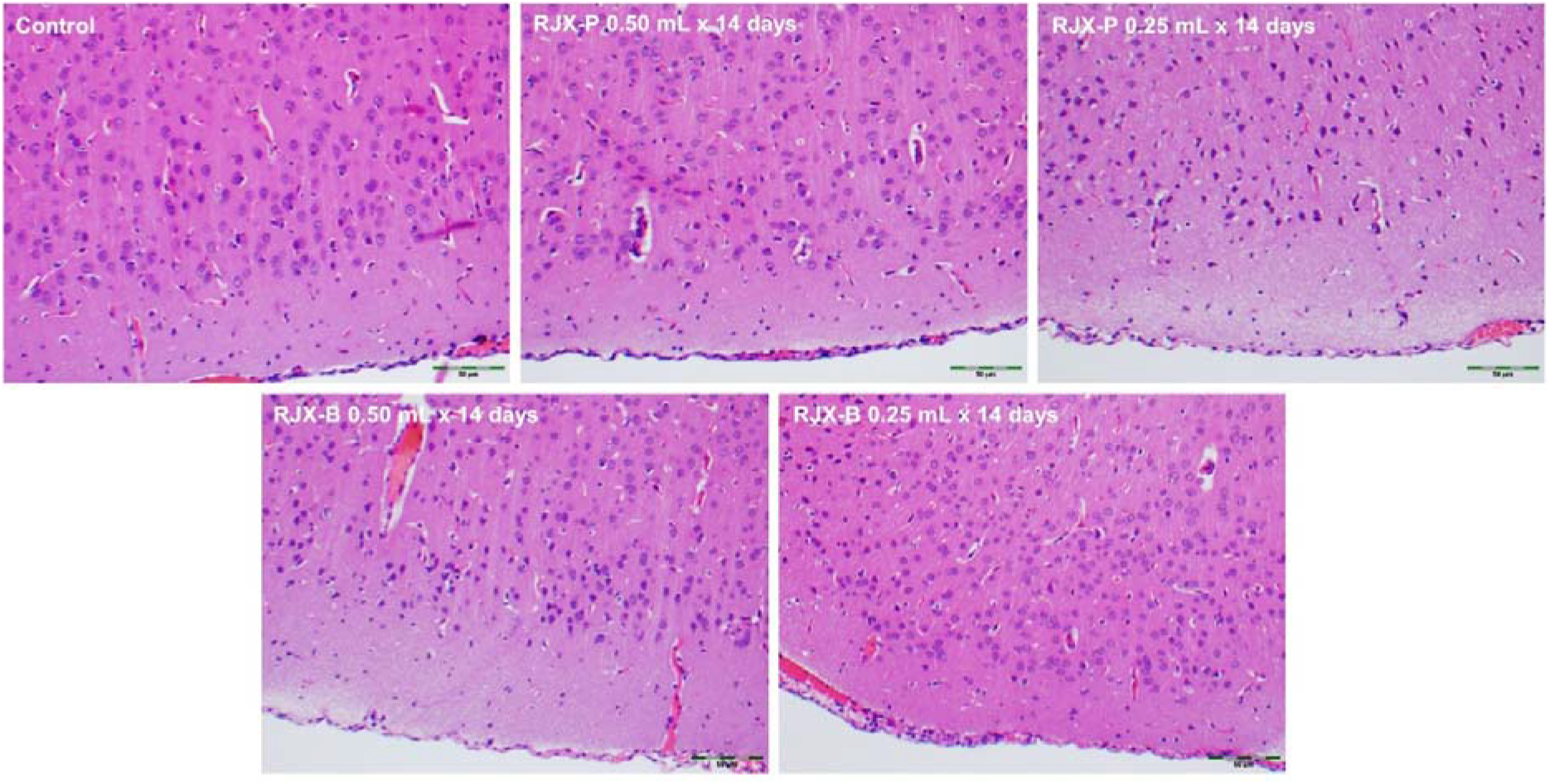
The day-15 histopathology of brain tissue in rats treated with 14 days of RJX-P or RJX-B intraperitoneal injections. Groups of 10 Wistar albino rats were treated with i.p injections of RJX-P (1.25 and 2.50 mL/kg/day) or RJX-B (1.25 and 2.50 mL/kg/day), or vehicle (NS) for 14 days. Animals were sacrificed on day 15 for histopathologic examinations. No toxic lesions were detected. H&E, 200x.

### Pharmacodynamics

14-day treatment with RJX-P and RJX-B similarly elevated the Day 15 tissue activities of the antioxidant enzyme superoxide dismutase (SOD) as well as ascorbic acid levels in both the lungs and liver in a dose-dependent fashion (**Table 11, Figure 7, Figure 8**). The activity of SOD and ascorbic acid were significantly higher in tissues of RJX-P or RJX-B treated rats than vehicle-treated control rats (*p*<0.0001). There was no statistically significant difference between SOD activity or ascorbic acid tissue levels of rats treated with RJX-P vs. rats treated with RJX-B (*p*>0.05). The observed elevations of the SOD activity and ascorbic acid levels were transient and were no longer detectable on day 28 following a 14-day recovery period (**Figure 9, Figure 10**). These results demonstrate that RJX-P and RJX-B are bioequivalent relative to their pharmacodynamic effects on tissue SOD activity and ascorbic acid levels.

**Table 11.**
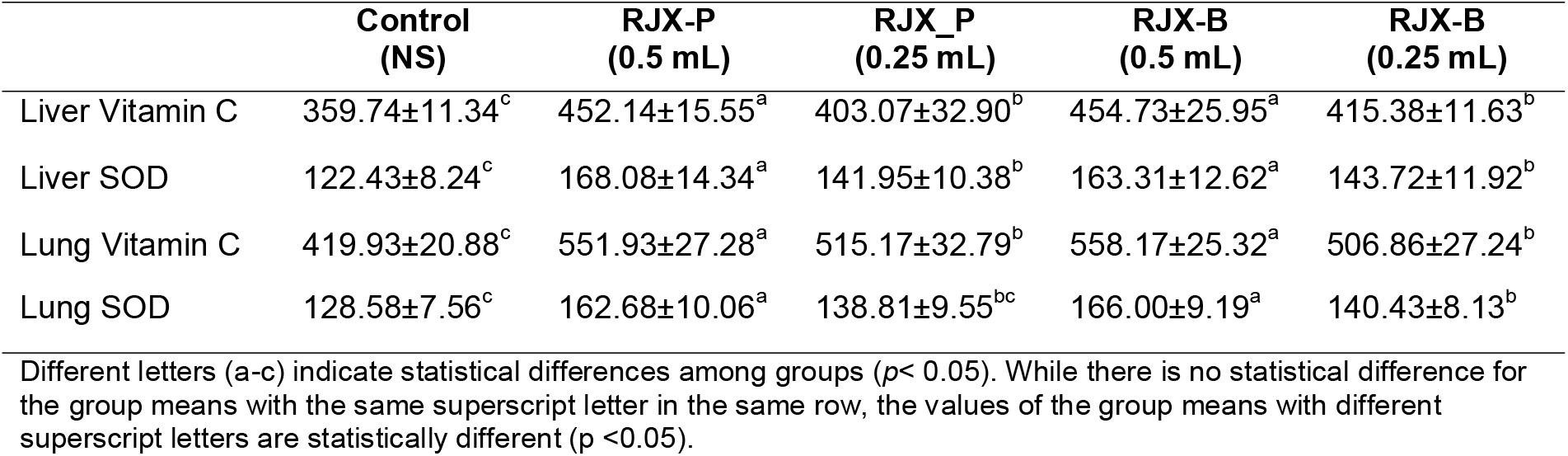
Comparison of the dose-dependent pharmacodynamic effects of 14-day treatment with RJX vs. RJX-B on liver and lung tissue SOD (U/mg protein) and ascorbic acid (μg/g) levels.

|  | Control<br>(NS) | RJX-P<br>(0.5 mL) | RJX_P<br>(0.25 mL) | RJX-B<br>(0.5 mL) | RJX-B<br>(0.25 mL) |
| --- | --- | --- | --- | --- | --- |
| Liver Vitamin C | 359.74±11.34<sup>c</sup> | 452.14±15.55<sup>a</sup> | 403.07±32.90<sup>b</sup> | 454.73±25.95<sup>a</sup> | 415.38±11.63<sup>b</sup> |
| Liver SOD | 122.43±8.24<sup>c</sup> | 168.08±14.34<sup>a</sup> | 141.95±10.38<sup>b</sup> | 163.31±12.62<sup>a</sup> | 143.72±11.92<sup>b</sup> |
| Lung Vitamin C | 419.93±20.88<sup>c</sup> | 551.93±27.28<sup>a</sup> | 515.17±32.79<sup>b</sup> | 558.17±25.32<sup>a</sup> | 506.86±27.24<sup>b</sup> |
| Lung SOD | 128.58±7.56<sup>c</sup> | 162.68±10.06<sup>a</sup> | 138.81±9.55<sup>bc</sup> | 166.00±9.19<sup>a</sup> | 140.43±8.13<sup>b</sup> |

Different letters (a-c) indicate statistical differences among groups ( $p < 0.05$ ). While there is no statistical difference for the group means with the same superscript letter in the same row, the values of the group means with different superscript letters are statistically different ( $p < 0.05$ ).

**Figure 7.**
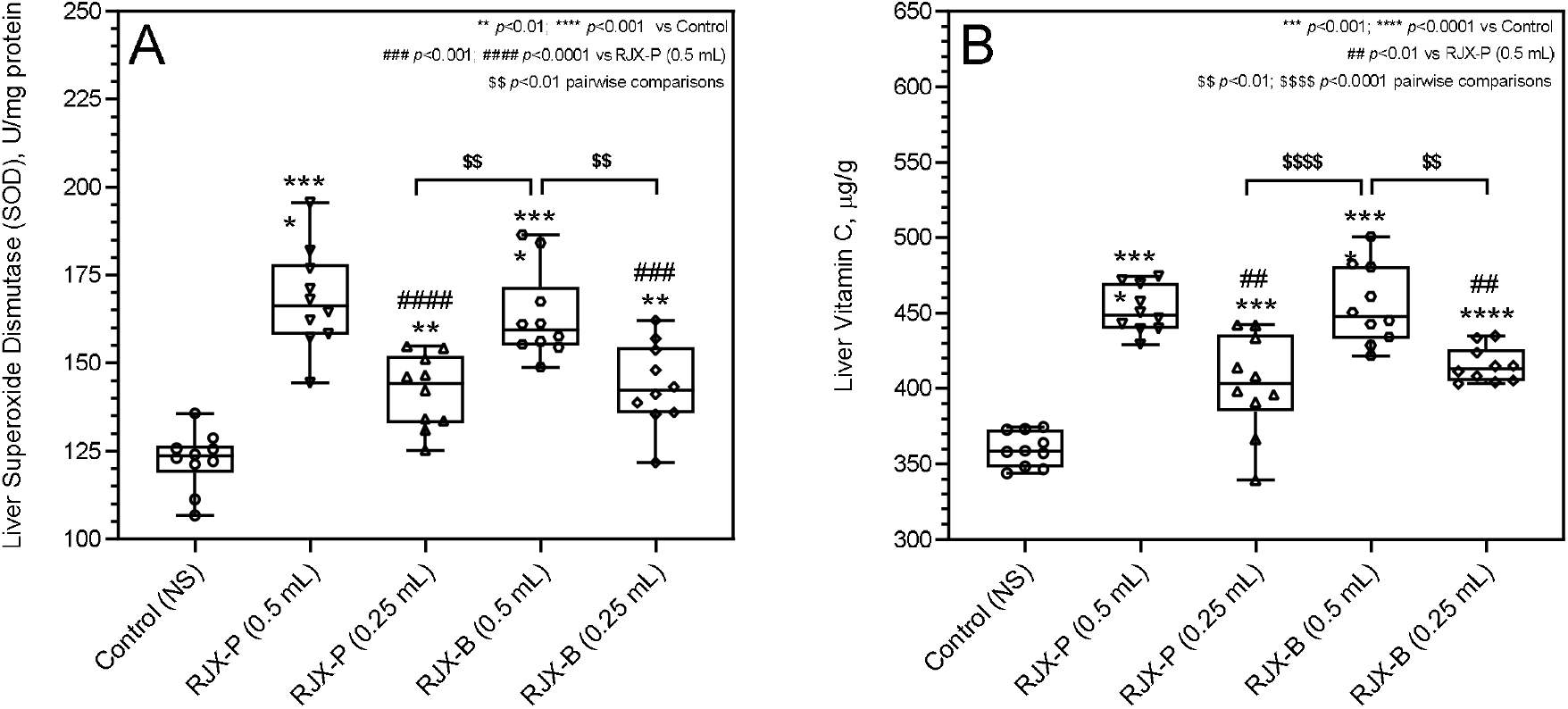
Comparison of the dose-dependent pharmacodynamic effects of 14-day treatment with RJX-P vs. RJX-B on liver tissue superoxide dismutase (SOD) and ascorbic acid levels. The depicted Whisker plots represent the median and values for the liver SOD (Panel A) and ascorbic acid (Panel B) levels. (ANOVA and Tukey’s post-hoc test were used for comparing the results among different treatment groups. Statistical significance between groups is shown by: ** p<0.01; *** p<0.001; p<0.0001 compared as Control group and, ## p<0.01; ### p<0.001; #### p<0.0001 compared as RJX-P (0.5 mL) group and, $$ p<0.01; $$$$ p<0.0001 pairwise comparisons between the groups).

**Figure 8.**
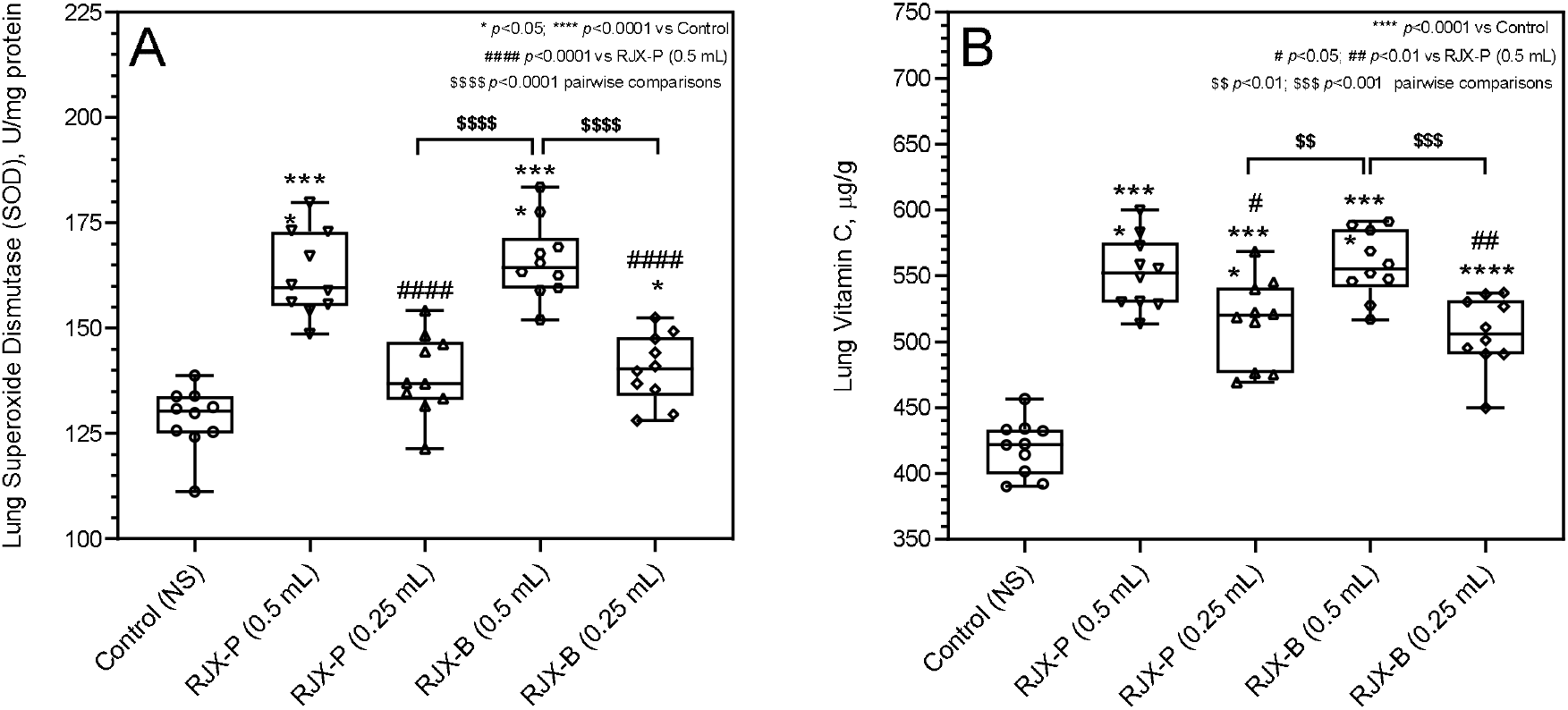
Comparison of the dose-dependent pharmacodynamic effects of 14-day treatment with RJX-P vs. RJX-B on lung tissue superoxide dismutase (SOD) and ascorbic acid levels. The depicted Whisker plots represent the median and values for the lung SOD (Panel A) and ascorbic acid (Panel B) levels (ANOVA and Tukey’s post-hoc test were used for comparing the results among different treatment groups. Statistical significance between groups is shown by: * p<0.05; **** p<0.0001 compared as Control group and, # p<0.05; ## p<0.01; #### p<0.0001 compared as RJX-P (0.5 mL) group and, $$ p<0.01; $$$$ p<0.0001 pairwise comparisons between the groups).

**Figure 9.**
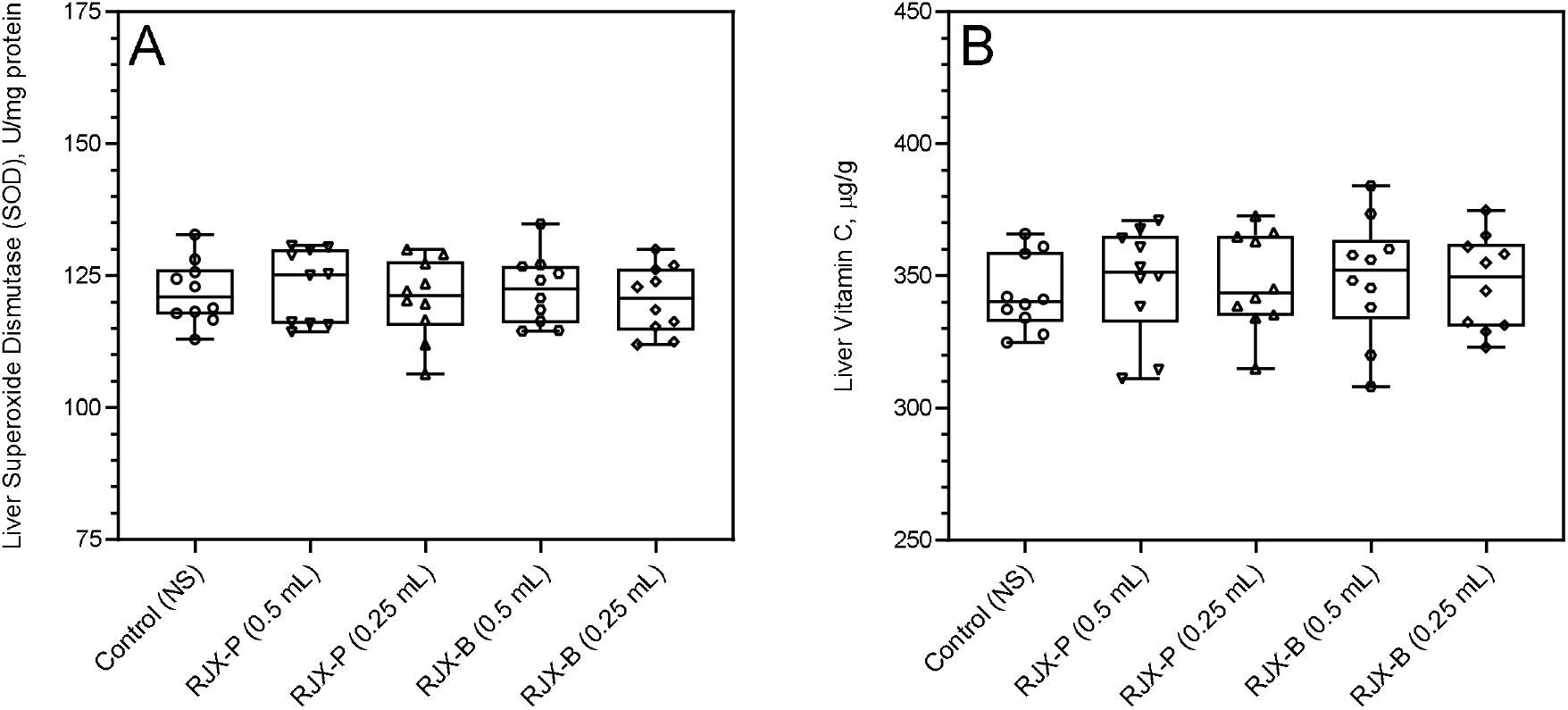
Comparison of the dose-dependent pharmacodynamic effects of 14-day treatment with RJX-P vs. RJX-B and sacrificed on day 28 following a 2 week recovery period on liver tissue superoxide dismutase (SOD) and ascorbic acid levels. The depicted Whisker plots represent the median and values for the liver SOD (Panel A) and ascorbic acid (Panel B) levels (ANOVA and Tukey’s post-hoc test were used for comparing the results among different treatment groups).

**Figure 10.**
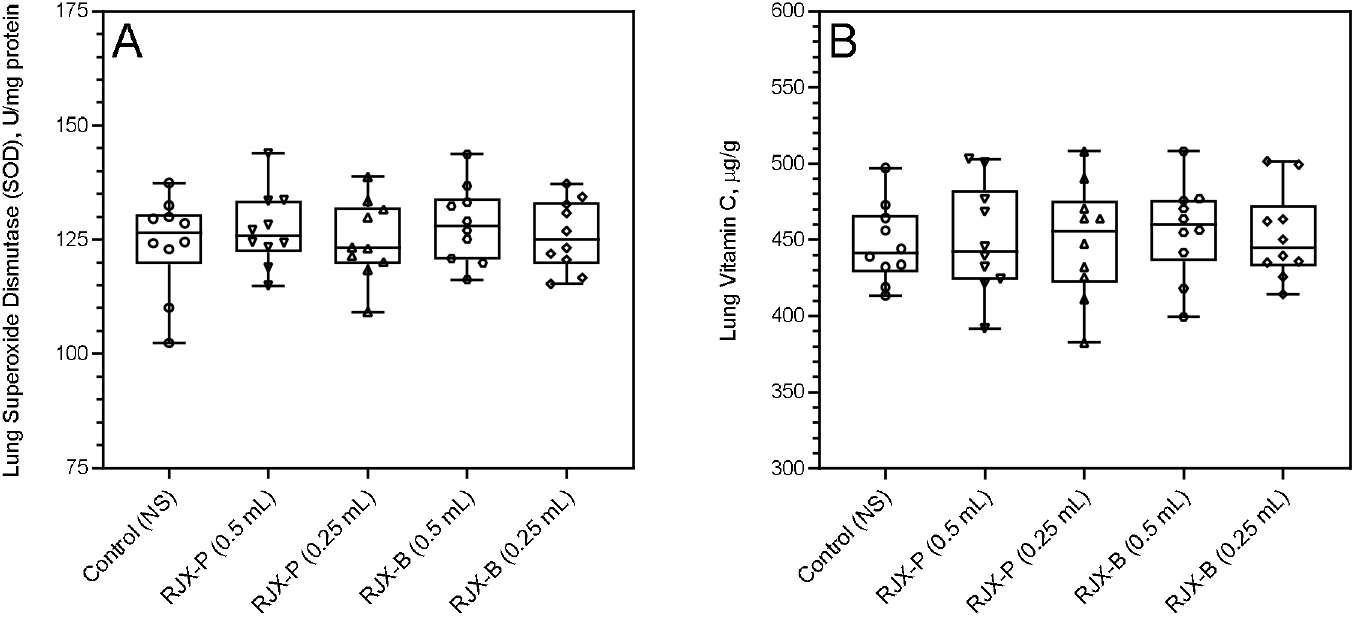
Comparison of the dose-dependent pharmacodynamic effects of 14-day treatment with RJX-P vs. RJX-B and sacrificed on day 28 following a 2 week recovery period on lung tissue superoxide dismutase (SOD) and ascorbic acid levels. The depicted Whisker plots represent the median and values for the lung SOD (Panel A) and ascorbic acid (Panel B) levels (ANOVA and Tukey’s post-hoc test were used for comparing the results among different treatment groups).

## Discussion

Patients with high-risk COVID-19 are in urgent need of effective treatment strategies that can prevent the development of ARDS and multi-organ failure [1–4]. Recently, we demonstrated that RJX prevents in a mouse model of sepsis the marked increase of each of proinflammatory cytokines like IL-6, TNF-α, and TGF-β in the serum as well as lungs and liver when treatments are initiated prophylactically prior to the injection of a fatal dose of LPS-GalN [3]. Notably, the activities of the antioxidant enzymes SOD, CAT, and GSH-Px were reduced after injection of LPS-GalN consistent with severe oxidative stress. RJX substantially improved the abnormally reduced activities of the antioxidant enzymes SOD, CAT, and GSH-Px, and levels of ascorbic acid. These results established RJX as a potent antioxidant with strong anti-inflammatory activity. The present study demonstrates that a 14-day treatment with RJX formulations increases tissue levels of ascorbic acid as well as SOD. RJX-P and RJX-B were bioequivalent relative to their pharmacodynamic effects on tissue SOD activity and ascorbic acid levels. These results confirm the anti-oxidant activity profile of RJX. RJX could favorably affect the clinical course of high-risk COVID-19 and reduce its case mortality rate. Both formulations of RJX are anticipated to shorten the time to resolution of ARDS by eliminating the contributions of pro-inflammatory cytokines of IL-6, TNF-α and TGF-β to the systemic and pulmonary inflammatory process as well as mitigating sepsis-associated oxidative stress.

## Author contributions

Each author (F.M.U., C.O., J.P., E.S., I.H.O., M.V., K.S.) has made significant and substantive contributions to the study, reviewed and revised the manuscript, provided final approval for submission of the final version. No medical writer was involved. F.M.U conceived the study, designed the evaluations reported in this paper, directed the data compilation and analysis, analyzed the data, and prepared the initial draft of the manuscript. Each author had access to the source data used in the analyses I.H.O. performed the necropsies and histopathologic examinations on mice.

## Funding/Support

This study was funded by Reven Pharmaceuticals, LLC, a wholly-owned subsidiary of Reven Holdings Inc.

## Conflicts of Interest

F.M.U., J.P. and M.V are employees/contractors of Reven Pharmaceuticals, the sponsor for the clinical development of RJX. C.O., E.S., I.H.O, and K.S declare no current competing financial interests.

## Role of the Funder/Sponsor

The sponsor did not participate in the collection of data. Three of the authors (F.M.U, J.P., and M.V) who participated in the analysis and decision to submit the manuscript for publication are affiliated with the sponsor.

